# Comprehensive analysis of the RBP regulome reveals functional modules and drug candidates in liver cancer

**DOI:** 10.1101/2024.09.04.611258

**Authors:** Mateusz Garbulowski, Riccardo Mosca, Carlos J. Gallardo-Dodd, Claudia Kutter, Erik L. L. Sonnhammer

## Abstract

RNA binding proteins (RBPs) are essential components of the transcriptomic regulome. Identifying the RBP regulome in cancer cells is crucial to discovering and understanding carcinogenesis mechanisms and providing new therapeutic targets. Here, we aimed to reveal the regulome of liver cancer upon specific perturbations. To this end, we applied a consensus Gene Regulatory Network (GRN) approach using knockdown data for the liver cancer cell line HepG2. By incorporating multiple GRNs from diverse inference methods, we constructed a highly precise GRN. To validate our results, we comprehensively evaluated the consensus GRN, focusing on characterizing the most relevant aspects of the liver cancer regulome. This included utilizing eCLIP-seq and RAPseq data to verify RBP interactions and binding sites. In addition, we performed an enrichment analysis of network modules and drug repurposing based on the inferred GRN. Taken together, our findings demonstrate the critical roles of RBP regulatory interactions in liver cancer that can be employed to improve treatment strategies.

## Introduction

RNA binding proteins (RBPs) are compounds that control diverse gene expression mechanisms. By involving a wide repertoire of post-transcriptional processes in cells, they support various biological functions such as decay, splicing, stabilization, translation, and transport (**Figure S1A**). RBPs bind RNA targets through a well-established RNA binding domain (RBD) referred to as canonical RBD. Studies on RBPs role in cancer imply that they participate in various mechanisms of progression, metastasis, and drug resistance (Gebauer et al. 2021; Pereira et al. 2017; Cen et al. 2023; King et al. 2011; Mucha et al. 2022; Qin et al. 2020). A recent study highlighted the role of RBPs in breast cancer drug resistance (Cen et al. 2023). Other work demonstrated that *CEBPZ*, characterized as RBP and transcription factor (TF), is associated with maintaining the leukaemic state (Barbieri et al. 2017). Yet another study has described the pro-tumorigenic role of a splicing factor in various cancers, *RBM39* (Xu et al. 2021). Furthermore, a set of prognosis-related RBPs have been obtained for liver cancer, among others including *EEF1E1*, *LIN28B,* and *XPO5* (Apizi et al. 2022). In addition to liver cancer-related RBPs, *PES1* affects the survival of liver cancer patients and enhances proliferation and tumorigenesis through the PI3K/AKT pathway (Wang et al. 2019). Therefore, it is of great interest to study interactions between RBPs as they often co-bind their targets (Quattrone and Dassi 2019). However, the RBP regulome in cancer is still poorly investigated; thus, exploring it is essential. This work focuses on the discovery of liver cancer regulome. Liver cancer is one of the top cancer-leading deaths that remains a therapeutic challenge (Llovet et al. 2022). There are multiple risk factors for liver cancer such as alcohol consumption, viral infections, obesity, diabetes, and other metabolic diseases (Ajoolabady et al. 2023). Although, nonalcoholic steatohepatitis is the fastest-growing cause of liver cancer (Huang et al. 2022). Previous studies investigated interactions among genes in liver cancer due to their importance in determining the treatment. For instance, it has been shown that the activation of *EGFR* reduces the response of liver cancer to the drug lenvatinib (Jin et al. 2021). Another example is that *LIN28B-AS1* and *IGF2BP1* bind to each other and promote liver cancer progression in human cells (Zhang et al. 2020). Moreover, it was depicted that oncogenesis in liver cancer is activated via the MYC signaling pathway (Dang et al. 2017).

Over the span of the last years, gene regulatory networks (GRNs) have been successfully engaged to analyze the mechanisms of various cancers (Xu et al. 2022; Madhamshettiwar et al. 2012; Emmert-Streib et al. 2014). For instance, GRN inference allowed for characterizing regulatory mechanisms of RBPs in pluripotency (Li and Izpisua Belmonte 2018). GRN estimates and displays interactions between regulators and their targets in biological systems using various omics data. In such graphic illustrations, nodes represent genes, and edges (or links) correspond to interactions between genes, the latter can be directed and signed. The common issue in the GRN field is noise in biological data that is often high and increases the risk of capturing false positive links. Thus, a consensus approach is applied to decrease the amount of false positives. Lately, a benchmarking study has concluded that knowledge about a perturbation profile is essential to infer an accurate GRN (Seçilmiş et al. 2022). Thus experiments that allow for controlled perturbation of gene expression such as shRNA knockdown followed by RNA-seq (shRNA-seq) shall be widely applied for GRN inference. Furthermore, it has been evaluated that the performance of the same GRN inference methods varies between diverse datasets (Shen et al. 2023). Due to such heterogeneity, an optimal solution to infer the most accurate GRN may not be definable. Therefore, an integrative approach should be investigated. The so-called consensus approach is inspired by the wisdom of crowds (WOC) principle where communities are more insightful than individuals. In other words, by integrating results from multiple perturbation-based GRN inference methods one can obtain better performance than using a single method (Marbach et al. 2012). To this end, several consensus approaches have been developed on biological data to support disease characterization and treatment (Meng et al. 2022; Chilingaryan et al. 2021). Moreover, several tools for the consensus GRN approach following the WOC idea have been created, for instance, based on a package for inferring consensus GRN, and by benchmarking, authors observed increased robustness and performance (Chilingaryan et al. 2021). Another example is GENECI which also exhibits high-quality GRN inference with a consensus approach (Segura-Ortiz et al. 2023).

In this study, we aimed to reveal the liver cancer regulome to provide RBP targets as candidates for further research and treatment. We achieved that with a consensus GRN approach by combining perturbation-based state-of-the-art regression and machine learning methods (Tjärnberg et al. 2017). The consensus GRN was constructed from the shRNA-seq ENCODE data created for HepG2 cells where a set of RBPs was considered. To support our results and methodology, we performed benchmarking of the consensus approach with synthetically generated shRNA-seq ENCODE-like data. The consensus GRN of liver cancer was validated comprehensively using in-house and public data resources, among others eCLIP-seq and RAPseq. Moreover, we estimated modules on liver cancer GRNs and performed module-specific gene enrichment analysis. To investigate protein-coding gene sets, we performed gene enrichment analysis on sets of common targets for RBP pairs. Finally, drug repurposing analysis was performed on the whole GRN as well as selected targets. Taken together, our comprehensive multi-step analysis allowed us to uncover the essential components of the liver cancer regulome.

## Results

### The regulome of liver cancer shows the dependency between *AQR* and *PES1*

To infer a reliable GRN of RBPs in liver cancer cells, we developed a consensus approach (**Supplementary text, Figure S1, S2, S3 and S4**) and applied it to the ENCODE shRNA-seq knockdown dataset. Complete consensus GRNs for HepG2 and K562 can be viewed in **Supplementary Table S1** and **S2**, respectively. We explored the liver cancer regulome using the 5+ consensus GRN for the HepG2 cell line, which contained 117 links. (**Figure 1A**). In this GRN, we observed three interactions of particular interest as they were detected by seven or more methods, indicating a high likelihood of being true positive links. These interactions are *AQR*-*PES1*, *RBM39*-*KIF1C,* and *FASTKD1*-*RPS10*. The 5+ methods threshold was chosen for the consensus GRN based on its high precision (0.70) in the benchmark, implying a high fraction of true positive links (**Figure S2 and S3**), while also maintaining a relatively high coverage and density, with an average outdegree of 1.65.

**Figure 1.**
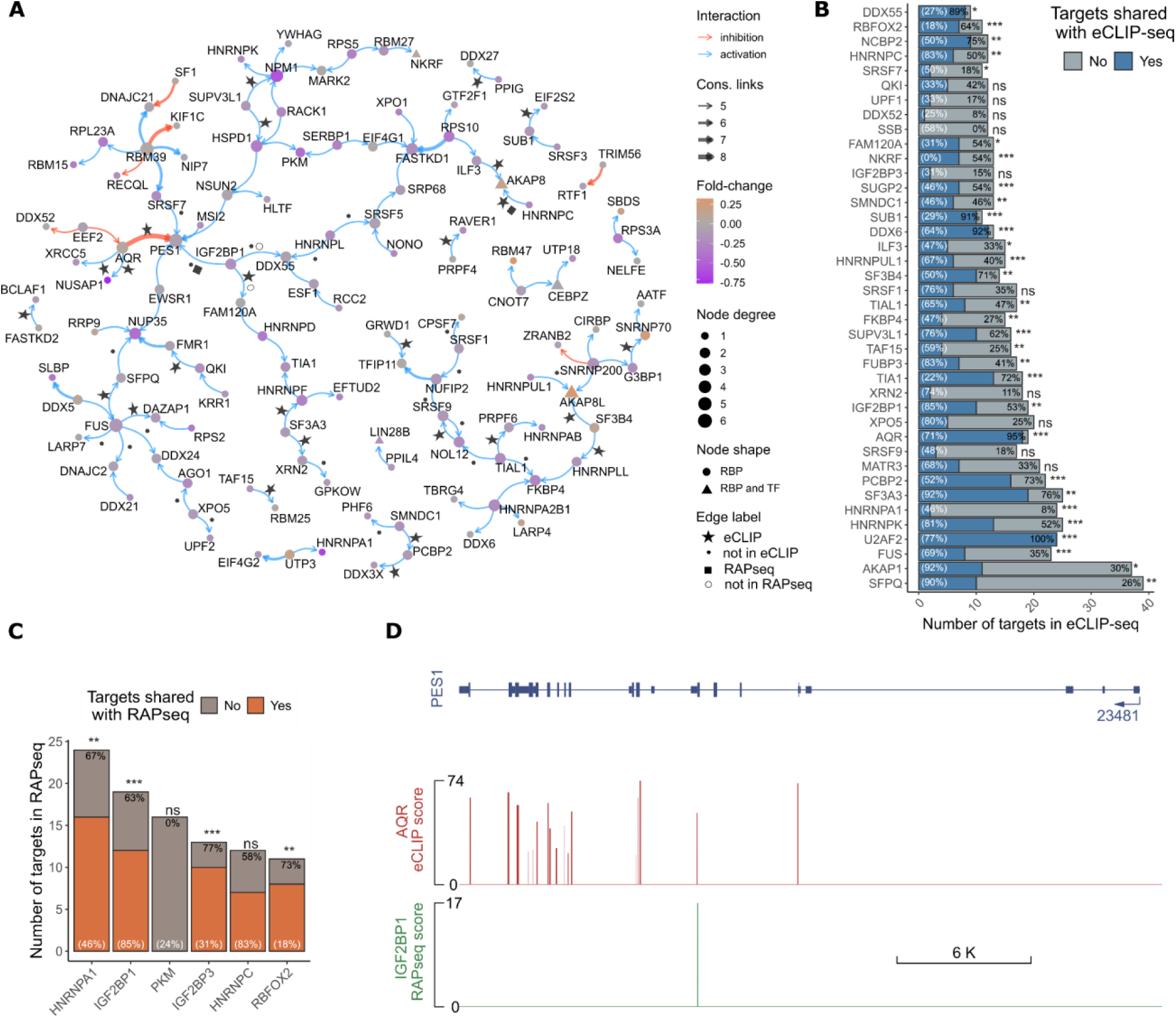
The regulome of liver cancer constructed as a consensus GRN **A.** Consensus GRN illustrating five minimum consensus links (5+). Edge width indicates the number of GRNs supporting the edge; node color indicates expression Log_2_ fold-change in perturbed HepG2 cells over control; and edge label indicates significant (FDR < 0.05) binding in eCLIP-seq or RAPseq data. **B-C.** Bargraphs show topmost hubs (outgoing node degree > 10) from the 2+ consensus GRN validated with eCLIP-seq and RAPseq data. Only RBPs that include eCLIP-seq or RAPseq data are shown in the plot. For each gene, a hypergeometric test was performed (not significant (ns): *p* ≥ 0.1; *: *p* = [0.05, 0.1]; **: *p* = [0.01, 0.05]; ***: *p* ≤ 0.01) to assess if a significant number of its targets are shared between the consensus GRN and significant (FDR < 0.05) experimental peaks. The shared links to targets are marked as the percentage of undirected RBP-RBP links in the GRN that overlap significant RBP-RBP interactions in eCLIP-seq or RAPseq. For each gene, its percentage of outgoing links in the GRN is given in white font. **D**. Genome track exemplifies AQR eCLIP-seq and IGF2BP1 RAPseq binding signals to *PES1* exonic (blue boxes) and intronic (blue line) region. Only significant (FDR < 0.05) peaks are shown. The number indicates gene ID.

We further analyzed the regulatory RBPs by using eCLIP-seq and RAPseq data and performing hypergeometric tests to evaluate the overlap between targets from undirected GRN and experimental data (**Figure 1A and 1B**). This analysis revealed the significant roles of *U2AF2* and *AQR* as major as well as *SUB1* and *DDX6* as minor RBP regulators in liver cancer, with over 90% of their targets confirmed (**Figure 1B**).

Notably, we confirmed the RBP regulation of *AQR*-*PES1* by the eCLIP-seq experiment. Specifically, we examined the binding sites in *PES1* when the expression of *AQR* was diminished (**Figure 1D**). This finding exemplifies that *PES1* is a target not only of *AQR* but also of other RBPs such as *IGF2BP1*. In the literature, it was described that *AQR*, encoding an RNA helicase, contributes to the accumulation of DNA damage upon knockdown in colon cancer cells (Sakasai et al. 2017). This suggests that *PES1* and *IGF2BP1* might be involved in this biological process in liver cancer as well. Notably, *PES1* has been investigated as a DNA damage-related factor in colorectal cancer (Xie et al. 2013). Overall, our findings underscore the importance of specific regulatory interactions among RBPs in liver cancer and highlight the potential roles of key RBPs, like AQR, PES1, and IGF2BP1 in biological processes.

### The validation of RBP interactions reveals their novelty and impact on survival

To reveal the regulome of liver cancer-related RBPs, we performed iterative enrichment analyses with DisGeNET as described in the methods section (see Validation of consensus GRN and **Figure S5**). We present the regulome (**Figure 2**) for the top 119 links, selected based on the highest local maximum and the lowest *p* value across all enrichments. In addition, we divided the GRN into two subGRNs corresponding to high and low FunCoup5 scores, reflecting either high or low FunCoup5 confidence scores of functional associations (**Figure 2A**, **2B**). The FunCoup5 evidence scores were discretized into low and high using equal frequency binning cutoffs. Both subGRNs can indicate pathway activation or complex formation in response to changes in a cell. Notably, in the low-scored subGRN, *IGF2BP1* emerged as a master regulator of other RBPs, confirmed with eCLIP-seq and RAPseq experiments. Moreover, we also detected interactions absent in the FunCoup5 database, marked with low* or high*, suggesting novel elements in the liver cancer regulome. A complete and signed liver cancer-related GRN is available in Supplementary Materials (**Figure S6A**). A full validation table for 3+ GRN is included in **Supplemenatry Table S3**. The information about applied databases can be found in **Supplementary Table S4**.

**Figure 2.**
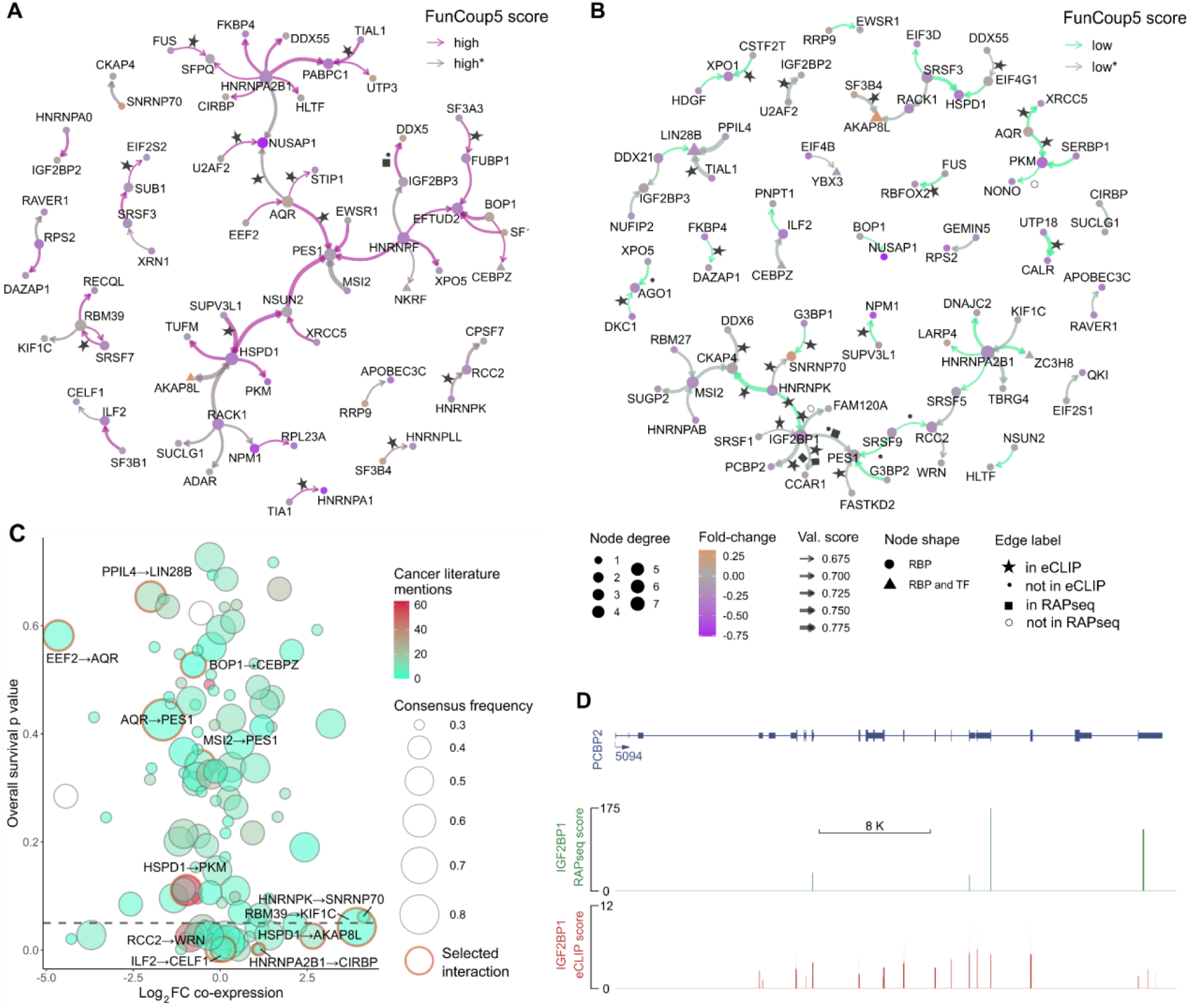
Liver cancer-related GRN and its validation. **A.** GRN of liver cancer-related RBPs with high FunCoup5 score. Edge width indicates the number of GRNs supporting the edge and edge colors indicate a high FunCoup5 score (bright violet labeled as high) and predicted high FunCoup5 score (faded violet labeled as high*). **B.** GRN of liver cancer-related RBPs with low FunCoup5 confidence score. Edge width indicates the number of GRNs supporting the edge and edge colors indicate a low FunCoup5 score (bright seagreen labeled as low) and predicted low FunCoup5 score (faded seagreen labeled as low*). **C.** Validation of the liver cancer regulome using the effect of RBP expression on patient overall survival, co-expression fold-change (FC), and RBP cancer literature popularity. Interaction pairs are shown with RBP names for some of the noteworthy predicted interactions listed in **Supplementary Table S5**. The dashed line indicates a *p* value of 0.05. **D.** Genome track exemplifies IGF2BP1 eCLIP-seq and RAPseq binding signals to *PCBP2* exonic (blue boxes) and intronic (blue line) region. Only significant (FDR < 0.05) peaks are shown. The number indicates gene ID.

The *IGF2BP1* is a notable RBP relevant to cancer pathways as previously investigated (Atanasoai et al. 2021; Müller et al. 2019). Given the availability of both eCLIP-seq and RAPseq data for *IGF2BP1*, we merged its significant targets from both experiments and performed enrichment analysis on such a set (**Figure S6B-D**). Our analysis revealed that *IGF2BP1* targets were associated with various disease states, including cancer and immune response. Its targets are also significantly related to Rho GTPases and the activation of the MYC pathway, as well as various transport and localization biological processes.

In the Supplementary Material, we provide a validation summary of all 119 liver cancer-related interactions (**Figure S7**) and a comprehensive validation list (**Supplementary Table S3)**. Selected interactions were further investigated (**Figure 2C**). For example, *HSPD1-AKAP8L* and *ILF2-CELF1* involve RBPs that strongly affect the survival of liver cancer patients, despite being understudied on cancer-related RBPs. We also observed a group of interactions that do not significantly affect survival (**Figure 2C**, points above the dashed line). Another example is *RBM39*-*KIF1C,* which exhibited high co-expression fold-changes and impact on survival but has low literature popularity. Conversely, *HSPD1*-*PKM* included RBPs that were highly mentioned in the literature concerning cancer. The oncogenic role of *HSPD1* has been investigated in oral and breast cancers (Kang et al. 2019; Kim et al. 2019a), while oncogenic activities of PKM were confirmed in thyroid and colorectal cancer (Desai et al. 2014; Kim et al. 2019b). In summary, multi-layer validation allowed us to confirm known and identify novel interactions between RBPs in liver cancer.

### Enrichment analysis of common protein coding targets of RBP interactions reveals activation of cancer-related pathways

In this section, we examined the complete regulome of 3+ GRN. We investigated all interactions to find overlapping targets between each pair of RBPs. Using targets derived from eCLIP-seq and RAPseq experiments, we analyzed RNA protein coding genes (**Figure S8**). As a result, we discovered three interactions that share the most significant number of RNA protein coding targets: *HNRNPK*-*HNRNPM*, *SFPQ*-*RBFOX2,* and *U2AF2*-*TIAL1*. Members of the heterogeneous nuclear ribonucleoprotein (hnRNP) family have been described as the main regulators of alternative splicing (Palombo et al. 2020). We identified that the *HNRNPK*-*HNRNPM* interaction had a high FunCoup5 score with a confidence level of 1 in the protein-protein interaction network. Enrichment analysis of their common targets indicated potential involvement in various cancer pathways, such as the Rho GTPases cycle, platelet-derived growth factors (PDGF), or VEGFA/VEGFR2 signaling (**Figure S8B**). We further found that *HNRNPM* has been proposed as a therapeutic target in liver cancer (Zhu et al. 2022). The *SFPQ*-*RBFOX2* interaction, with a FunCoup5 evidence score of 0.48 for complex formation, showed enrichmentpatterns similar to *HNRNPK*-*HNRNPM*, but also included the EGF/EGFR signaling pathway. Both genes belong to a group of splicing factors (Ivanova et al. 2023). Lastly, the *U2AF2*-*TIAL1* interaction has a FunCoup5 score of 0.766 for metabolic coupling. Except for several cancer-related pathways, such as ROBO or VEGFA/VEGFR2, their targets are associated with several immune response pathways (Whisenant 2017). The role of *U2AF2* in immune response and the importance of *TIAL1’s* function in adaptive immunity was covered by other works (Osma-Garcia et al. 2023; Whisenant 2017). These findings underscore the complex regulatory interactions involving RBPs and their potential roles in cancer and immune response pathways.

### Validation of transcription factors shows liver cancer-specific regulation among CEBPZ and RBM27

In the set of analyzed RBPs, several of them were also annotated as transcription factors (TFs). To perform a validation study of TFs, we used resources such as GRAND and GRNdb that included precomputed GRNs for liver cancer and healthy cells. In detail, we intersected the 3+ consensus GRN with the GRNs from GRAND and GRNdb, using undirected GRN to investigate all possible interactions.

Our analysis revealed a high relationship between *SFPQ* and *CEBPZ* across 24 liver cancer cell lines (**Figure S9**). However, it is also reflected in the healthy liver GRNs (**Figure S9D**). Another prominent interaction is *CEBPZ*-*RBM27*, which was absent in a healthy liver, but present in all investigated GRNs of liver cancer cells. Overall, the *CEBPZ* TF appears to regulate numerous RBPs, including *RBM22*, *ESF1,* or *ABCF1*. It has been shown that *CEBPZ*, also known as *CTF2*, physically interacts with *TP53* (Uramoto et al. 2003). *RBM27* is not frequently mentioned in the context of cancer regulation, but its role in RNA decay has been discovered (Silla et al. 2020). These findings underscore the pivotal roles of transcription factors like CEBPZ in regulating the expression of RBP genes in a cancer cell-specific manner.

### Module detection reveals groups of RBPs related to specific cancer pathways and MYC targets

To perform a module enrichment analysis, we utilized two GRNs, namely the 5+ consensus GRN (**Figure 1A**) and the 3+ consensus liver cancer-related GRNs (**Figure 2A-B, Figure S6A**). We observed that several modules contained RBPs related to the oncogenic transcription factor *MYC* (**Table 1**). It was established that *MYC* plays a crucial role in liver cancer initiation and progression (Sequera et al. 2022). Notably, the light green module (**Figure 3A**) includes four out of six RBPs (*NPM1, RACK1, HSPD1* and *SUPV3L1*) that were identified as *MYC* targets. This suggests that the remaining two RBPs, *YWHAG* and *HNRNPK,* could potentially be novel *MYC* targets. Members of the YWHA and HNRNP families have previously been identified as *MYC* targets (Leal et al. 2016; David et al. 2010). Within the same module, two RBP *MYC* targets from the MsigDB gene set HALLMARK_MYC_TARGETS_V2, *NPM1* activation by *SUPV3L1*, are connected. Their interaction was confirmed by eCLIP-seq, suggesting liver cancer-relevant regulation occurring in the *MYC* pathway. To validate these, we investigated co-expression mechanisms between *MYC* and the potentially novel *MYC* targets (**Figure S11**).

**Figure 3.**
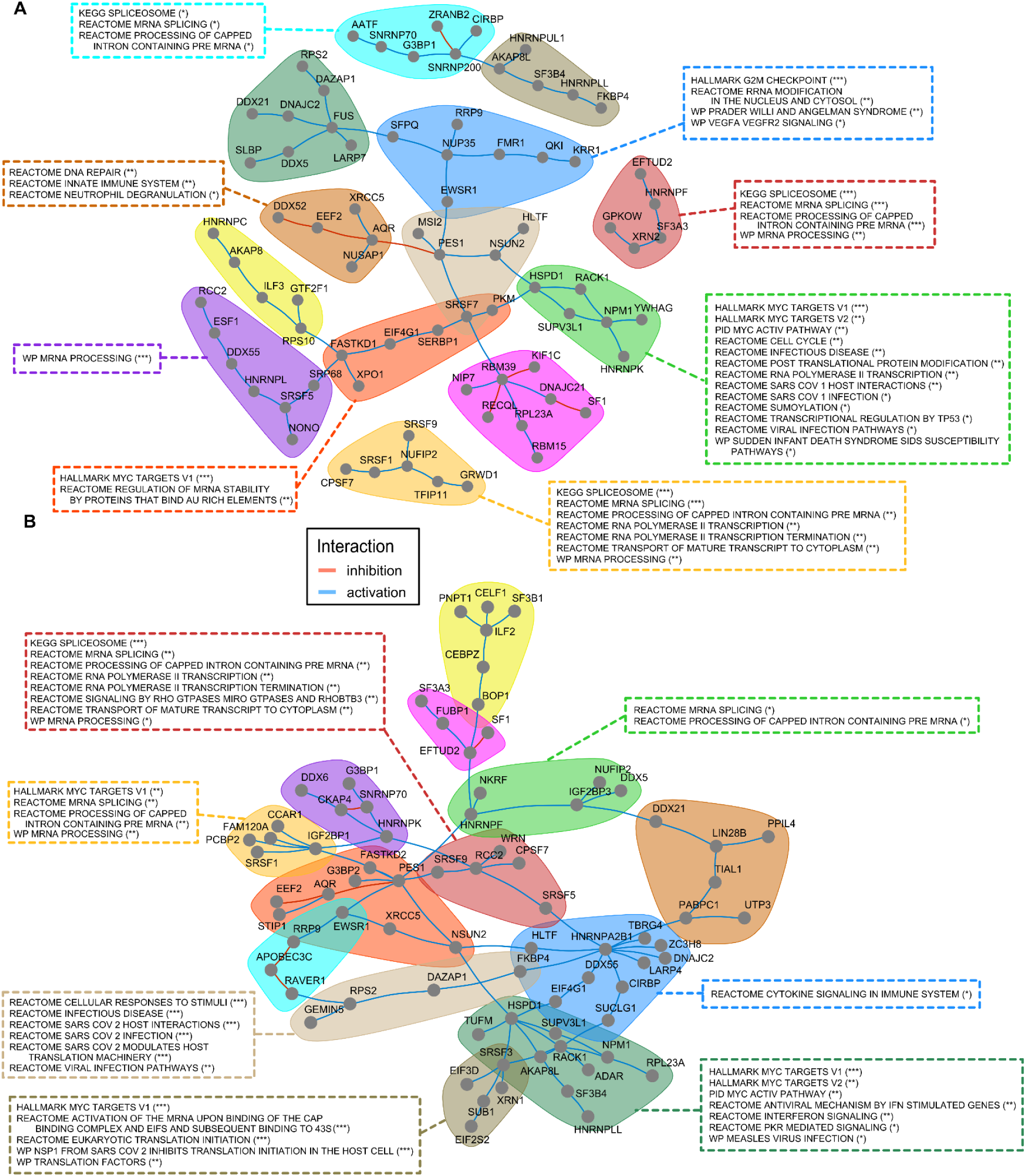
Module detection and MSigDB (C2, C5, C6, and H) enrichment of modules. Selected and simplified significant (FDR < 0.05) terms are included in the legend. As a background, a full list of RBPs was taken. Some terms were merged and simplified according to their similarity. **A.** The consensus GRN where at least 5 methods support each link. Modules are color-coded and pathway enrichment terms are displayed. **B.** GRN of 119 liver cancer-related links. Terms include FDR-corrected *p*-values as follows *p* ≥ 0.1; *: *p* = [0.05, 0.1]; **: *p* = [0.01, 0.05]; ***: *p* ≤ 0.01.

**Table 1.**
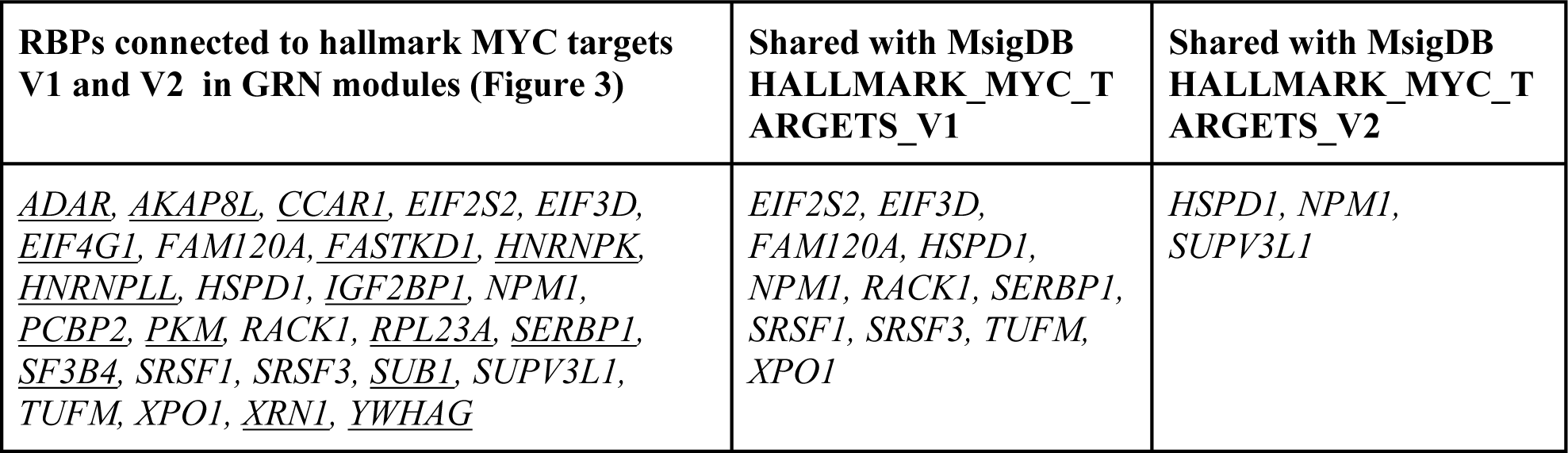
Hallmark *MYC* targets and connected RBPs found in GRN modules. Novel *MYC*-related genes are underlined.

Moreover, we identified clusters related to various cancer pathways. These included VEGFA/VEGFR2, Rho GTPases and DNA repair. For instance, the *AQR*-*XRCC5* interaction included in the light brown module (**Figure 3A**) contained RBPs involved in DNA repair pathways. Another noteworthy module is that of *IGF2BP1* (**Figure 3B**, light orange module), which was enriched with RBPs involved in mRNA splicing and processing of capped intron among other pathways (Rawat and Heemers 2024).

Several modules were also enriched with splicing-related terms, indicating the crucial role of splicing performed by RBPs during oncogenesis (Zhang et al. 2021). We found distinct splicing modules (and in **Figure 3A** dark red module and **Figure 3B** light orange clusters). Another significant enrichment was found for the immune system and infectious disease pathways in multiple clusters. This finding aligns with the growing recognition that immunotherapy is a promising strategy for treating liver cancer patients and improving their survival (Li et al. 2021).

Overall, our results provided an overview of pathways affected in liver cancer cells upon RBP knockdown and revealed potentially novel *MYC* targets that warrant further investigation for their carcinogenic potential.

### GRN-based drug repurposing identifies RBP-targeting drug candidates

To perform drug repurposing analysis on the liver cancer regulome, we performed a few analyses. First, we investigated the entire consensus 5+ GRN in CLUEreg, which revealed Irinotecan as the topmost drug based on cosine similarity. This suggested that HepG2 cells undergoing RBP knockdown may exhibit similar mechanisms to those induced by Irinotecan (**Table S6**), a well-known anticancer drug used for colorectal cancer treatment as it is an inhibitor of DNA topoisomerase I (Bailly 2019).

Second, we evaluated *IGF2BP1,* in line with a previous study that investigated its role in liver cancer (Atanasoai et al. 2021). We analyzed common eCLIP-seq and RAPseq-based targets of *IGF2BP1* (**Figure 2, Figure S6B, C, D**). Our analysis identified Garcinol or Gallic-acid as potential agents to reverse the expression profile of IGF2BP1. Garcinol inhibits STAT3, and Gallic-acid arrest cells at the G2/M phase. STAT3 has been shown to play a crucial role in cancer inflammation and immunity, aligning with the results of our enrichment analysis (**Figure S6C, D**). In addition, based on the consensus 2+ GRN, we evaluated the differentially expressed gene targets *AQR* and *U2AF2* (**Figure 1B**). Notably, the effect of targets of two major regulators *AQR* and *U2AF2* may be reversed by WNT-related drug treatments (**Table S6**) (He and Tang 2020; López-Pérez et al. 2023).

Finally, based on CTD, we found several chemotherapeutic drugs, such as Doxorubicin and Ivermectin (**Figure 4, Figure S12**). Recent studies have confirmed the role of these drugs in liver cancer therapy (Kim et al. 2017; Lu et al. 2022). Our analysis indicates that these two drugs contribute to inhibiting or activating RBP MYC targets. For example, Doxorubicin increases the expression level of *IGF2BP1,* which is very low in liver cancer patients, while Ivermectin and Acetaminophen decrease the expression of *IGF2BP1(Lu et al. 2022; Wu et al. 2024)*. Moreover, both Doxorubicin and Ivermectin decrease the expression of *PES1,* which is elevated in liver cancer patients (**Figure S12**). We also found phosphorylation-related interactions between caffeine and MYC-related targets, with multiple studies suggesting that coffee consumption may decrease the risk of various cancers, including liver cancer (Safe et al. 2023; Pauwels and Volterrani 2021).

**Figure 4.**
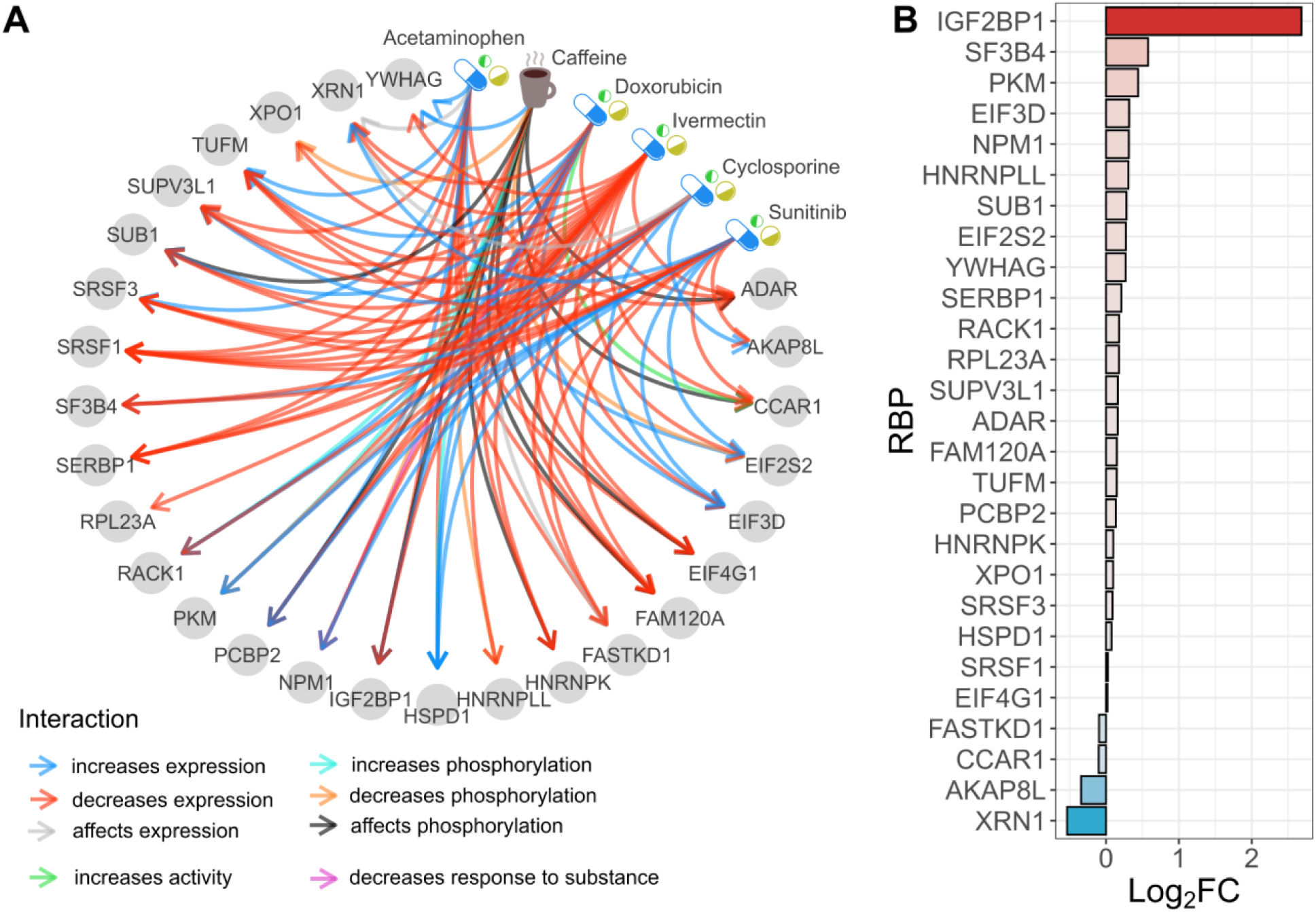
Drug repurposing and patients-based expression of *MYC*-related RBP targets (**Table 1**). **A.** CTD-derived interactions between drugs and RBPs. The color of the edges represents various types of interactions. Drugs are marked as nodes with special icons. **B.** Log_2_ fold-change (Log_2_FC) based on expression from TCGA and GTEx cohorts. The colors of the bars represent the negative (blue) or positive (red) Log_2_FC. Values were calculated as median RBP expression in LIHC over median RBP expression in healthy liver.

In sum, our findings highlight several promising drug candidates and potential therapeutic targets for liver cancer, providing new options for treatment strategies based on modulating RBPs and their regulatory network.

## Discussion

Mechanisms behind cancerogenesis are complex and often involve interactions among molecules that reflect the activation or deactivation of certain pathways. Multiple studies attempted to estimate regulatory networks for liver cancer, yet not focused on RBPs and often utilized only a single inference method (Meier et al. 2021; Zeng et al. 2012; Bakr et al. 2023). The RBP regulome of liver cancer represents crucial processes that may lead to activations and inhibitions in human cancer cells. To design treatment it is valuable to recognize these processes and provide hypotheses of binding mechanisms that influence the expression of RBPs and their targets. Furthermore, it is important to handle noise in gene expression data that can lead to false positive interactions. To produce a regulome in a more accurate and interpretable way, a consensus GRN approach can be applied. To this end, we utilized 10 various GRN inference methods to reveal the liver cancer regulome by following the WOC approach (Marbach et al. 2012) and further performed a comprehensive analysis.

A substantial outcome of our work is a set of interactions among 232 RBPs in liver cancer. Based on benchmarking with the ENCODE-like data, we believe that several RBP-RBP regulations are very likely to be true positives, namely *AQR*-*PES1*, *RBM39*-*KIF1C,* and *FASTKD1*-*RPS10*. Notably, *AQR*-*PES1* was detected by 8 out of 10 methods, and the eCLIP-seq experiment further confirmed the binding of these two RBPs. The co-expression in *AQR*-*PES1* is much lower for LIHC than GTEx (with a high FunCoup5 score), meaning that interaction between them is much weaker in liver cancer cells.

Another finding is the set of *MYC*-related RBP targets detected based on the module enrichment analysis of GRNs. As RBPs share a lot of common mechanisms related to posttranscriptional processes, it was not a trivial task to perform enrichment. Thus, we used a full list of known RBPs as a background. The enrichment results agree with the previous work (Liu et al. 2023) and show the important role of RBPs in the MYC pathway. We can also observe that some clusters (**Figure 3**) correspond to other cancer-related pathways, e.g. DNA repair, immune system response and splicing (Gillman et al. 2021; Batel et al. 2024). Regarding the drug repurposing analysis, we found several drug candidates previously reported as inhibitors of pathways that are related to cancer. We can observe that the drug candidates are linked to the induction of apoptosis and other major cancer pathways. For instance, Sunitinib is an inhibitor of *SF3B4* and an activator of *XRN1* that is upregulated and downregulated in liver cancer, respectively (**Figure 4**). This drug is a tyrosine kinase receptor inhibitor used for cancer treatment (Faivre et al. 2007).

As a result of simulations, we share two synthetic ENCODE-like data, together with corresponding GRNs, so that they can be reused in future studies for benchmarking these cohorts (see Data Availability). We also share full consensus GRNs and a 3+ consensus validation table that can be a resource for future comparison of regulomes (**Table S1-3**). The K562 cell line was also used in this work, however not analyzed as extensively as HepG2. This data and its consensus GRN are attached in supplementary materials to this work (**Table S2**). We believe that it can be comprehensively utilized in the future as a resource for interactions for studies on leukemia.

In this work, we aimed to create a consensus approach using only a perturbation-based method, thus the commonly applied GENIE3 was not considered. Being aware of the good performance of GENIE3 in benchmarks and other analyses, we used CART in the consensus approach. Furthermore, GENIE3 and its successor GRNBoost2 (Moerman et al. 2019) consider all unperturbed genes in data, while here we focus only on perturbed, i.e. knocked-down, RBPs that allow us to compute a consensus GRN with better performance. As our analysis is limited to GeneSPIDER-embedded methods and single-omics ENCODE data, a more complex approach utilizing a bigger collection of methods and multi-omics data for a consensus approach could be investigated in the future. For example, by involving methods that rely on the information about transcription factors. This could result in discovering even more true positives while keeping a denser consensus GRN.

While analyzing our data, we can see a limitation to the perturbed set of RBPs, thus any other targets, e.g. protein coding genes, were not investigated for inferring GRNs. However, this work focused on interactions among RBPs as they play a crucial role in cells (see Introduction). Regarding non-RBP genes, based on eCLIP-seq and RAPseq cohorts, we performed enrichment analysis of targets that included protein coding genes (**Figure S6 and S8**). This allowed us to find pathways of protein coding targets that are controlled by RBPs. Moreover, the eCLIP-seq and RAPseq collection is another limitation as not all 232 RBPs were validated. Future studies could perform RBP-RNA binding experiments to corroborate interactions, e.g. for *RBM39* that exhibited regulatory role in this study, for which ENCODE eCLIP-seq is missing.

In conclusion, the analysis allowed us to reveal the RBP-related regulome of liver cancer together with its functional modules and drug candidates. To the best of our knowledge, it is the first study that infers a liver cancer regulome of RBPs via a consensus approach using perturbation-based methods. The discovery of liver cancer regulome relies on comprehensive benchmarking that reflects the high precision of GRNs using a consensus approach. Our findings brought known and novel interactions among RBPs. For the inferred consensus GRN, we performed comprehensive validation using external data sets and GRNs. Furthermore, we executed a GRN-based module enrichment analysis that revealed a set of known and potentially novel *MYC* targets. Our analysis detected several RBP-RBP interactions likely to participate in cancer-related pathways in the liver tissue. For instance, *AQR*-*PES1* is the strongest evidence in this analysis. Finally, we performed drug repurposing that revealed a list of potential cancer drug candidates. Overall, the results from module enrichment analysis and drug repurposing can support the development of treatment for liver cancer patients.

## Methods

This section includes a description of the data and methods used for the analysis. We also include a summary picture of our research (see **Figure 5**).

**Figure 5.**
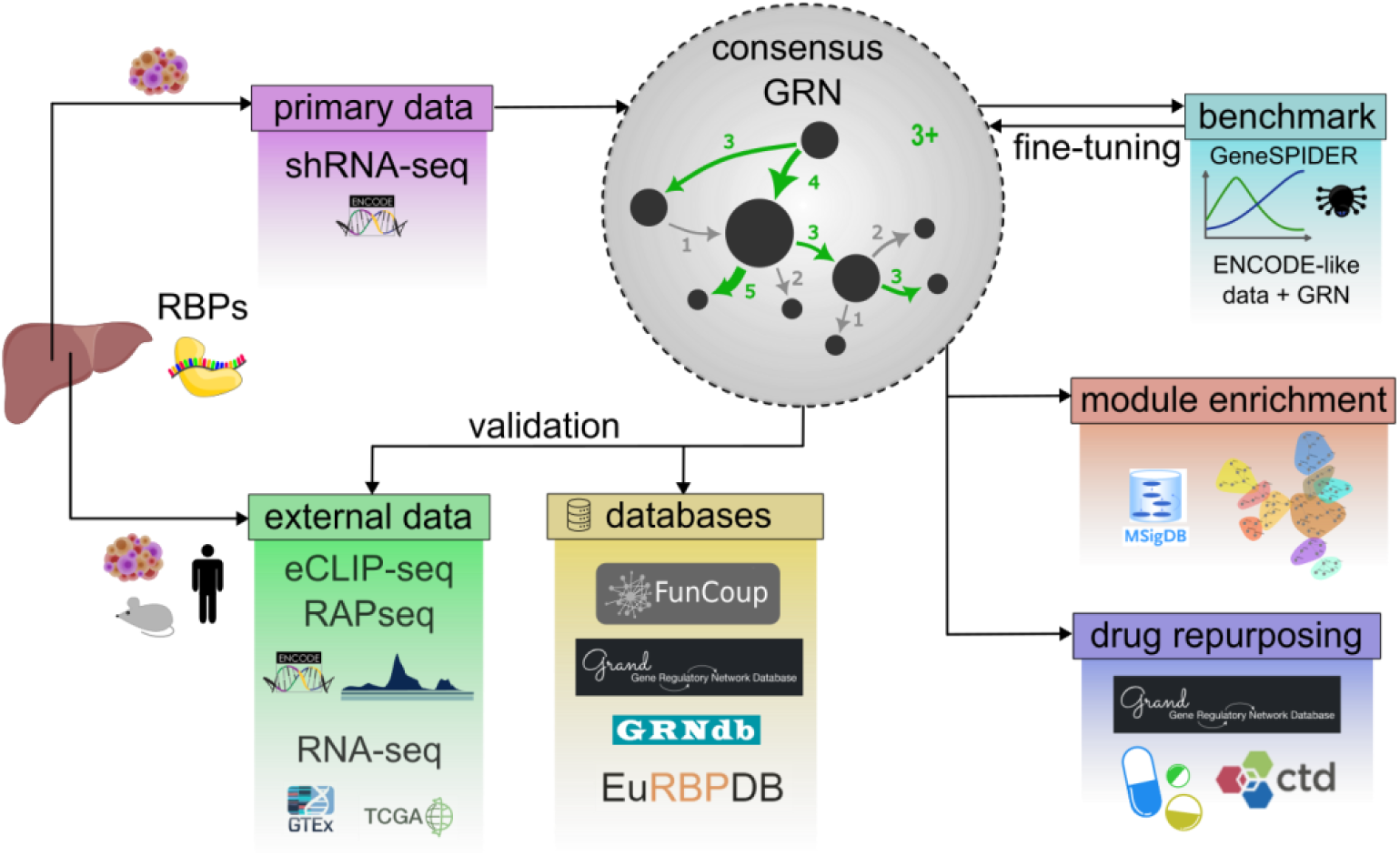
An overview of the analysis aiming at elucidating liver cancer mechanisms between RBPs.

### shRNA knockdown followed by RNA-seq

The GRN inference was conducted on publicly available shRNA RNA-seq of the liver cancer cell line (HepG2), provided by the Encyclopedia of DNA Elements (ENCODE) project (Luo et al. 2020). The ENCODE repository aims at generating and disseminating data on functional elements of the human genome. In addition, we incorporated shRNA RNA-seq from the chronic myelogenous leukemia cell line (K562) to generate a corresponding GRN for comparison, i.e. a control GRN. In both ENCODE data sets, cells were treated with shRNA targeting 232 RBPs. To infer GRNs from these data sets, we separated the replicates within samples. This allowed us to adhere to the perturbation design necessary for the employed GRN inference methodology (Tjärnberg et al. 2017). The final datasets comprised perturbed genes, referenced against the GRCh38 genome, presented as log_2_ fold-change expression matrices.

### Binding sites detection with eCLIP-seq and RAPseq

To analyze and validate the RBP regulome, we utilized data for enhanced crosslinking and immunoprecipitation sequencing (eCLIP-seq) (Van Nostrand et al. 2016) from ENCODE and RNA affinity purification followed by sequencing (RAPseq) from the in-house repository (Atanasoai et al. 2021). Both methods enable transcriptome-wide profiling of RBP binding sites, albeit with procedural differences. The main difference between these two techniques is their experimental protocol (Atanasoai et al. 2021) since eCLIP-seq is performed *in cellulo* while RAPseq is conducted *in vitro*. The number of profiled RBPs is much larger for eCLIP-seq (n=103) than for RAPseq (n=9) experiments considered RBP-RNA interactions significant for peaks with a *p*-value (P) < 0.05. In total 106 RBP-RNA interactomes from the combined eCLIP-seq and RAPseq datasets were used for validation.

### ENCODE-based synthetic data and benchmarking

To generate ENCODE-like synthetic datasets supplied with gold standard GRNs, we utilized GeneSPIDER (Tjärnberg et al. 2017) aiming at inference and benchmarking with controlled GRN and data properties (publicly available at bitbucket.org/sonnhammergrni/genespider). In the simulation, we assumed the same data properties as the real ENCODE shRNA RNA-seq, i.e. data size of 232 RBPs and two replicates. In addition, based on **Figure S1B**, GRN was generated with the average outgoing node degree of 3 (excluding the self-loops). To find the synthetic data set that is the most similar to ENCODE, we iteratively simulated gene expression data with various signal-to-noise ratios (SNRs) and GRNs. In the first step, we started by drawing *SNR*_*L* between 0.0001 and 0.1 and calculated the Pearson correlation coefficient (Matlab function *corrcoef*) among replicate pairs. The SNR is defined as:

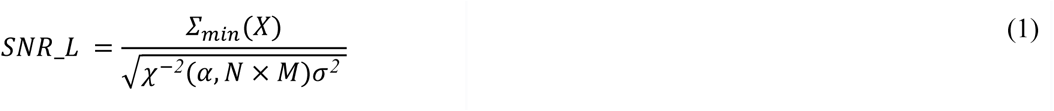

where *∑*_*min*_(*X*) is the smallest singular value of the noise-free gene expression matrix *X*, *χ^−2^(α, N × M)* is the inverse *χ^2^*distribution at level *α* with *N* × *M* degrees of freedom (number of genes × number of experiments), and *σ^2^* is the variance of the additive noise matrix. The synthetic gene expression was generated as steady state data (*Y*) following the linear mapping: *Y* = −*A^−1^P + E*, where *A* is an adjacency matrix of a GRN, with negative real part for eigenvalues, *P* is the matrix of the perturbation design, and *E* is the matrix of Gaussian noise defined at the *SNR*_*L* level. To mimic the ENCODE data *P* is a diagonal matrix where −1 indicates the knockdown of a given gene and 0 indicates unperturbed genes. Next, we compared average correlation *μ*_*ρ*_*E*_ of simulated data to the average correlation *μ*_*ρ*_*S*_ of the real ENCODE data as *μ*_*ρ*_*diff*_ = |*μ*_*ρ*_*E*_ − *μ*_*ρ*_*S*_|. The procedure was executed 1000 times for 20 randomly generated scale-free GRNs. The corresponding GRN and data with the lowest *μ*_*ρ*_*diff*_ were kept as the closest to the real data. Specifically, we selected synthetic data for HepG2 and K562 with *μ*_*ρ*_*diff*_ = 1.62e-05 and *μ*_*ρ*_*diff*_ = 1.73e-04, respectively. This corresponded to *SNR*_*L*=0.0055 for HepG2 and *SNR*_*L*=0.0029 for K562. The synthetic ENCODE-like datasets generated in this manner, along with their corresponding gold standard GRNs, were used for benchmarking.

The ENCODE-like data was used to perform benchmarking on the consensus GRN approach and tune the methodology towards the real ENCODE data. To assess the performance of the consensus-based approach and investigate how many true positive links are kept, we used a positive predictive value (*PPV*) measure:

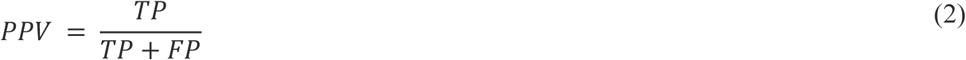

where TP is the number of true positive links and FP is the number of false positive links. We also used more global performance metrics such as the F1 score, which is the harmonic mean of precision and recall:

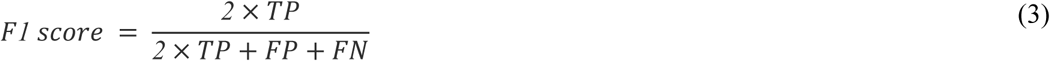

where FN is the number of false negative links.

### Inference of consensus GRNs

To infer a consensus GRN, we employed the following 10 methods: normalized least squares (LSCON) (Hillerton et al. 2022), ridge regression with least square cutoff (ridgeco), total least squares with cut-off (tlsco), lasso, logistic regression with lasso (loglasso), classification with support vector machine (svmc), z-score (Zscore), decision trees (CART), regression neural networks (neunetreg) and linear gaussian processes (gaplin) included in the GeneSPIDER framework (Tjärnberg et al. 2017). Most of these methods have been efficiently used in previous studies and analyses (Seçilmiş et al. 2022; Morgan et al. 2020; Seçilmiş et al. 2020). Recently introduced methods, including decision trees and neural network regression, have also been successfully applied for the GRN inference (Huynh-Thu et al. 2010; Moerman et al. 2019; Shu et al. 2021). However, we benchmarked and excluded two other methods. Namely, least squares, due to its tendency to overestimate the GRN, and the elastic net, as it performed slightly worse than lasso while obtaining almost identical GRN.

In constructing the consensus GRN, we employed the WOC approach by averaging the number of links detected across all inferred GRNs (**Figure S1C**). Each link was assigned a weight based on the number of methods that inferred it, divided by the total number of methods used. For example, a link with a weight of 0.5 means that it was inferred by five methods, and a GRN with 5+ consensus links includes only those links detected by at least five methods. In this work, we also used a 3+ GRN, which had a precision of around 0.5 and included 648 interactions. The 3+ GRN served as a larger resource for validation and to retrieve true positive links not present in the 5+ GRN. Moreover, in some instances, we also used a 2+ GRN to validate targets of RBP hubs.

The sign of the links, indicating inhibition or activation, in the consensus GRN was established by using the WOC approach as well. The sign was based on inference methods capable of sign estimation, i.e. all methods except CART and neunetreg. In addition, we also used Spearman correlation coefficients (*ρ*) calculated for each interaction using the ENCODE expression data, where replicates were averaged. For indecisive cases, for instance, when two methods gave negative signs and two others inferred positive signs, the sign was taken based on the *ρ*.

### Validation of consensus GRN

To validate the 3+ consensus GRN, we used several resources and external data sets. These included: 1) gene expression from 110 healthy liver tissue samples from GTEx, 2) gene expression from 421 samples from Liver Hepatocellular Carcinoma (LIHC) samples from TCGA, and 3) clinical data with survival statistics of LIHC from TCGA. To perform a fair comparison, unified UCSC Xena cohorts with transcripts per million (TPM) gene expression values were used for the former mentioned three datasets (Goldman et al. 2017). GTEx and TCGA cohorts were employed to calculate Pearson’s correlation coefficient of LIHC and healthy liver tissue. The values given in the validation tables represent the difference in the coefficients between LIHC and healthy liver tissue. Additionally, 4) from EuRBPDB, we incorporated literature mentions of RBPs related to cancers and differential expression of RBPs detected in case-control studies of cancer into a “cancer literature popularity score” (Liao et al. 2020). 5) we also used ENCODE-derived eCLIP-seq data for HepG2 and K562 cells, representing a landscape of RBPs-RNA interactions, annotated with GENCODE v45 (Frankish et al. 2021) using the *queryGff* R function from the RCAS package (v1.22.0). These eCLIP-seq data were merged with significant peaks from the RAPseq dataset (22). 6) An in-house RBP list and an estimated number of literature mentions of genes being RBPs (Søndergaard et al. 2022) were also included. Furthermore, 7) we included functional association networks from FunCoup5, aimed at detecting direct and indirect functional associations based on regulatory mechanisms or pathways, constructed upon 10 evidence types, including genomics, proteomics, and transcriptomics data (Persson et al. 2021). Here, discrete association scores absent in FunCoup5 were predicted using decision trees, with a cross-validation accuracy of 85%, applied to tabular data including all the other validation features (points 1-6) used as a training set. Decision trees were chosen for their efficiency in learning from tabular data, fast performance, handling of missing data, and the learning process is interpretable (Song and Lu 2015). Training and testing were performed using the rpart package (v4.1.19).

To obtain a comprehensive table including all categorical features, non-categorical variables were clustered using parameterized finite Gaussian mixture models (GMM) (Mclust function from the mclust v6.1 R package) into three groups. In the case of having multiple zero values in a variable, GMM was applied to non-zero values to obtain two groups, and a zero value was taken as the third group. This approach was applied for variables including co-expression difference, literature-related features, and signal values from eCLIP-seq and RAPseq. Other features, such as survival significance group, differential expression significance group, consensus frequency, and presence in the K562 GRN and RBD group were treated as categorical variables. Next, all features were scaled between 0 and 1. To estimate the total score, we calculated the average of scores for each interaction and sorted the table in descending order. In the final step, the table was subdivided by incrementally selecting the top interactions in a top-down way and enriching RBPs included in the selected set of interactions. This enrichment was performed with the *enrichDGN* function using the DOSE package (v3.22.1). We selected terms and their *p* values where “liver” or “hepato” phrases were present. We then adjusted *p* values for False Discovery Rate (FDR) within each run and provided a single FDR value calculated as a harmonic mean of all terms (R package harmonicmeanp v3.0.1). In the final step, we selected a list of interactions giving the lowest average FDR.

### Validation of TFs with external GRNs

In a separate part of our analysis, we analyzed a set of TFs included as a part of our data. We incorporated several bulk and single-cell RNA-seq-based GRNs from GRAND and GRNdb (Ben Guebila et al. 2022; Fang et al. 2021) From GRAND, we collected liver cancer-related GRNs corresponding to 24 different cell lines and one set of TCGA liver-cancer patients, estimated with LIONESS and OTTER, respectively (Ben Guebila et al. 2023). From GRNdb, we used TF GRNs of human liver-related cancers derived from single-cell data (Fang et al. 2021). In GRNdb, GRNs were inferred using GENIE3 (Huynh-Thu et al. 2010). The GRNs from GRAND and GRNdb were intersected with our 3+ consensus GRN.

### Module detection and enrichment analysis

To detect modules on GRNs, we applied the igraph (1.4.1) function *cluster_infomap* that minimizes the expected description length of a random walker trajectory (Csárdi et al. 2024) as benchmarks showed their good performance (Choobdar et al. 2019). Initially, we identified clusters in GRN to filter out disconnected nodes. Subsequently, modules containing five or more genes were retained for further analysis. Then, we reapplied the detection of modules on the filtered GRN using the same approach. This allowed us to get much cleaner GRN and larger modules for the enrichment analysis. Next, we employed the clusterProfiler (4.4.4) package and its *enricher* function to assess the significance of the overlap between modules and the MSigDB gene sets (Subramanian et al. 2005; Liberzon et al. 2015). Specifically, we selected curated (C2) and hallmark (H) gene sets, using an in-house RBP list as a background (Søndergaard et al. 2022). Afterward, we collected *p* values for each module and processed the results by 1) removing terms enriched with only one gene, 2) performing module-oriented FDR adjustment, and 3) keeping only terms with significant FDR values (FDR < 0.05).

### Drug repurposing on the selected targets

To identify potential drug candidates for liver cancer treatment, we performed drug repurposing through three approaches. First, we submitted the entire signed 5+ consensus GRN to the CLUEreg tool (Ben Guebila et al. 2022). Second, we provided target sets for master regulators to CLUEreg. We selected all targets of IGF2BP1 based on eCLIP-seq and RAPseq, further limited to include the top 20% of differentially expressed genes (DEGs) based on Student’s t-test using TCGA and GTEx liver-related cohorts. Additionally, based on **Figure 1B**, we investigated all targets of AQR and U2AF2 based on 2+ consensus GRN, similarly limited to the top 20% of DEGs. As CLUEreg requires lists of down- and up-regulated targets as low- and high-targeted genes respectively, we utilized the aforementioned information about DEGs. The drugs were selected based on their high cosine similarity and low tau from the top 100 estimated by CLUEreg.

The third approach involved intersecting lists of selected RBPs with the Comparative Toxicogenomics Database (CTD) (Davis et al. 2021). Specifically, we used CTD to identify *MYC*-related targets identified through module enrichment analysis (**Table 1**) and a list of the most noteworthy interactions (**Table S5**). The output of CTD was then intersected with DrugBank to compile a list of drugs (Knox et al. 2024). Finally, we retained drugs that interacted with 50% or more of the RBPs in the specified set.

## Availability of data and materials

Simulated ENCODE-like data are publicly available at https://zenodo.org/records/12165429. The HepG2 shRNA-seq is publicly available via the ENCODE repository at https://www.encodeproject.org/biosamples/ENCBS282XVK/. The GeneSPIDER package is publicly available at https://bitbucket.org/sonnhammergrni/genespider/.

## Acknowledgments

We would like to thank Davide Buzzao, Dimitri Guala, and Thomas Hillerton for meaningful discussions.

## Funding

This work was supported by a postdoc grant from the Science for Life Laboratory’s SFO program [M.G.]. Funding for open access charge: Stockholm University.

## Notes

### Competing Interest Statement

The authors have declared no competing interest.

## References

Ajoolabady A, Tang D, Kroemer G, Ren J. 2023. Ferroptosis in hepatocellular carcinoma: mechanisms and targeted therapy. Br J Cancer 128: 190–205.

Apizi A, Wang L, Wusiman L, Song E, Han Y, Jia T, Zhang W. 2022. Establishment and verification of a prognostic model of liver cancer by RNA-binding proteins based on the TCGA database. Transl Cancer Res 11: 1925–1937.

Atanasoai I, Papavasileiou S, Preiß N, Kutter C. 2021. Large-scale identification of RBP-RNA interactions by RAPseq refines essentials of post-transcriptional gene regulation. bioRxiv 2021.11.08.467743. https://www.biorxiv.org/content/biorxiv/early/2021/11/09/2021.11.08.467743 (Accessed April 10, 2024).

Bailly C. 2019. Irinotecan: 25 years of cancer treatment. Pharmacol Res 148: 104398.

Bakr S, Brennan K, Mukherjee P, Argemi J, Hernaez M, Gevaert O. 2023. Identifying key multifunctional components shared by critical cancer and normal liver pathways via SparseGMM. Cell Rep Methods 3: 100392.

Barbieri I, Tzelepis K, Pandolfini L, Shi J, Millán-Zambrano G, Robson SC, Aspris D, Migliori V, Bannister AJ, Han N, et al. 2017. Promoter-bound METTL3 maintains myeloid leukaemia by m6A-dependent translation control. Nature 552: 126–131.

Batel A, Polović M, Glumac M, Šuman O, Jadrijević S, Lozić B, Petrović M, Samardžija B, Bradshaw NJ, Skube K, et al. 2024. Correction: SPRTN is involved in hepatocellular carcinoma development through the ER stress response. Cancer Gene Ther. 10.1038/s41417-024-00772-w.

Ben Guebila M, Lopes-Ramos CM, Weighill D, Sonawane AR, Burkholz R, Shamsaei B, Platig J, Glass K, Kuijjer ML, Quackenbush J. 2022. GRAND: a database of gene regulatory network models across human conditions. Nucleic Acids Res 50: D610–D621.

Ben Guebila M, Wang T, Lopes-Ramos CM, Fanfani V, Weighill D, Burkholz R, Schlauch D, Paulson JN, Altenbuchinger M, Shutta KH, et al. 2023. The Network Zoo: a multilingual package for the inference and analysis of gene regulatory networks. Genome Biol 24: 45.

Cen Y, Chen L, Liu Z, Lin Q, Fang X, Yao H, Gong C. 2023. Novel roles of RNA-binding proteins in drug resistance of breast cancer: from molecular biology to targeting therapeutics. Cell Death Discov 9: 52.

Chilingaryan G, Abelyan N, Sargsyan A, Nazaryan K, Serobian A, Zakaryan H. 2021. Combination of consensus and ensemble docking strategies for the discovery of human dihydroorotate dehydrogenase inhibitors. Sci Rep 11: 11417.

Choobdar S, Ahsen ME, Crawford J, Tomasoni M, Fang T, Lamparter D, Lin J, Hescott B, Hu X, Mercer J, et al. 2019. Assessment of network module identification across complex diseases. Nat Methods 16: 843–852.

Csárdi G, Nepusz T, Müller K, Horvát S, Traag V, Zanini F, Noom D. 2024. igraph for R: R interface of the igraph library for graph theory and network analysis. Zenodo https://zenodo.org/doi/10.5281/zenodo.7682609.

Dang H, Takai A, Forgues M, Pomyen Y, Mou H, Xue W, Ray D, Ha KCH, Morris QD, Hughes TR, et al. 2017. Oncogenic Activation of the RNA Binding Protein NELFE and MYC Signaling in Hepatocellular Carcinoma. Cancer Cell 32: 101–114.e8.

David CJ, Chen M, Assanah M, Canoll P, Manley JL. 2010. HnRNP proteins controlled by c-Myc deregulate pyruvate kinase mRNA splicing in cancer. Nature 463: 364–368.

Davis AP, Grondin CJ, Johnson RJ, Sciaky D, Wiegers J, Wiegers TC, Mattingly CJ. 2021. Comparative Toxicogenomics Database (CTD): update 2021. Nucleic Acids Res 49: D1138–D1143.

Desai S, Ding M, Wang B, Lu Z, Zhao Q, Shaw K, Yung WKA, Weinstein JN, Tan M, Yao J. 2014. Tissue-specific isoform switch and DNA hypomethylation of the pyruvate kinase PKM gene in human cancers. Oncotarget 5: 8202–8210.

Emmert-Streib F, de Matos Simoes R, Mullan P, Haibe-Kains B, Dehmer M. 2014. The gene regulatory network for breast cancer: integrated regulatory landscape of cancer hallmarks. Front Genet 5: 15.

Faivre S, Demetri G, Sargent W, Raymond E. 2007. Molecular basis for sunitinib efficacy and future clinical development. Nat Rev Drug Discov 6: 734–745.

Fang L, Li Y, Ma L, Xu Q, Tan F, Chen G. 2021. GRNdb: decoding the gene regulatory networks in diverse human and mouse conditions. Nucleic Acids Res 49: D97–D103.

Frankish A, Diekhans M, Jungreis I, Lagarde J, Loveland JE, Mudge JM, Sisu C, Wright JC, Armstrong J, Barnes I, et al. 2021. GENCODE 2021. Nucleic Acids Res 49: D916–D923.

Gebauer F, Schwarzl T, Valcárcel J, Hentze MW. 2021. RNA-binding proteins in human genetic disease. Nat Rev Genet 22: 185–198.

Gillman R, Lopes Floro K, Wankell M, Hebbard L. 2021. The role of DNA damage and repair in liver cancer. Biochim Biophys Acta Rev Cancer 1875: 188493.

Goldman M, Craft B, Zhu J, Haussler D. 2017. Abstract 2584: The UCSC Xena system for cancer genomics data visualization and interpretation. Cancer Res 77: 2584–2584.

He S, Tang S. 2020. WNT/β-catenin signaling in the development of liver cancers. Biomed Pharmacother 132: 110851.

Hillerton T, Seçilmiş D, Nelander S, Sonnhammer ELL. 2022. Fast and accurate gene regulatory network inference by normalized least squares regression. Bioinformatics 38: 2263–2268.

Huang DQ, Singal AG, Kono Y, Tan DJH, El-Serag HB, Loomba R. 2022. Changing global epidemiology of liver cancer from 2010 to 2019: NASH is the fastest growing cause of liver cancer. Cell Metab 34: 969–977.e2.

Huynh-Thu VA, Irrthum A, Wehenkel L, Geurts P. 2010. Inferring regulatory networks from expression data using tree-based methods. PLoS One 5. 10.1371/journal.pone.0012776.

Ivanova OM, Anufrieva KS, Kazakova AN, Malyants IK, Shnaider PV, Lukina MM, Shender VO. 2023. Non-canonical functions of spliceosome components in cancer progression. Cell Death Dis 14: 77.

Jin H, Shi Y, Lv Y, Yuan S, Ramirez CFA, Lieftink C, Wang L, Wang S, Wang C, Dias MH, et al. 2021. EGFR activation limits the response of liver cancer to lenvatinib. Nature 595: 730–734.

Kang B-H, Shu C-W, Chao J-K, Lee C-H, Fu T-Y, Liou H-H, Ger L-P, Liu P-F. 2019. HSPD1 repressed E-cadherin expression to promote cell invasion and migration for poor prognosis in oral squamous cell carcinoma. Sci Rep 9: 8932.

Kim DW, Talati C, Kim R. 2017. Hepatocellular carcinoma (HCC): beyond sorafenib-chemotherapy. J Gastrointest Oncol 8: 256–265.

Kim S-K, Kim K, Ryu J-W, Ryu T-Y, Lim JH, Oh J-H, Min J-K, Jung C-R, Hamamoto R, Son M-Y, et al. 2019a. The novel prognostic marker, EHMT2, is involved in cell proliferation via HSPD1 regulation in breast cancer. Int J Oncol 54: 65–76.

Kim Y, Lee Y-S, Kang SW, Kim S, Kim T-Y, Lee S-H, Hwang SW, Kim J, Kim EN, Ju J-S, et al. 2019b. Loss of PKM2 in Lgr5+ intestinal stem cells promotes colitis-associated colorectal cancer. Sci Rep 9: 6212.

King CE, Cuatrecasas M, Castells A, Sepulveda AR, Lee J-S, Rustgi AK. 2011. LIN28B promotes colon cancer progression and metastasis. Cancer Res 71: 4260–4268.

Knox C, Wilson M, Klinger CM, Franklin M, Oler E, Wilson A, Pon A, Cox J, Chin NEL, Strawbridge SA, et al. 2024. DrugBank 6.0: the DrugBank Knowledgebase for 2024. Nucleic Acids Res 52: D1265–D1275.

Leal MF, Ribeiro HF, Rey JA, Pinto GR, Smith MC, Moreira-Nunes CA, Assumpção PP, Lamarão LM, Calcagno DQ, Montenegro RC, et al. 2016. YWHAE silencing induces cell proliferation, invasion and migration through the up-regulation of CDC25B and MYC in gastric cancer cells: new insights about YWHAE role in the tumor development and metastasis process. Oncotarget 7: 85393–85410.

Liao J-Y, Yang B, Zhang Y-C, Wang X-J, Ye Y, Peng J-W, Yang Z-Z, He J-H, Zhang Y, Hu K, et al. 2020. EuRBPDB: a comprehensive resource for annotation, functional and oncological investigation of eukaryotic RNA binding proteins (RBPs). Nucleic Acids Res 48: D307–D313.

Liberzon A, Birger C, Thorvaldsdóttir H, Ghandi M, Mesirov JP, Tamayo P. 2015. The Molecular Signatures Database (MSigDB) hallmark gene set collection. Cell Syst 1: 417–425.

Li M, Izpisua Belmonte JC. 2018. Deconstructing the pluripotency gene regulatory network. Nat Cell Biol 20: 382–392.

Liu F, Liao Z, Zhang Z. 2023. MYC in liver cancer: mechanisms and targeted therapy opportunities. Oncogene 42: 3303–3318.

Li X, Ramadori P, Pfister D, Seehawer M, Zender L, Heikenwalder M. 2021. The immunological and metabolic landscape in primary and metastatic liver cancer. Nat Rev Cancer 21: 541–557.

Llovet JM, Castet F, Heikenwalder M, Maini MK, Mazzaferro V, Pinato DJ, Pikarsky E, Zhu AX, Finn RS. 2022. Immunotherapies for hepatocellular carcinoma. Nat Rev Clin Oncol 19: 151–172.

López-Pérez A, Remeseiro S, Hörnblad A. 2023. Diet-induced rewiring of the Wnt gene regulatory network connects aberrant splicing to fatty liver and liver cancer in DIAMOND mice. Sci Rep 13: 18666.

Lu H, Zhou L, Zuo H, Le W, Hu J, Zhang T, Li M, Yuan Y. 2022. Ivermectin synergizes sorafenib in hepatocellular carcinoma via targeting multiple oncogenic pathways. Pharmacol Res Perspect 10: e00954.

Luo Y, Hitz BC, Gabdank I, Hilton JA, Kagda MS, Lam B, Myers Z, Sud P, Jou J, Lin K, et al. 2020. New developments on the Encyclopedia of DNA Elements (ENCODE) data portal. Nucleic Acids Res 48: D882–D889.

Madhamshettiwar PB, Maetschke SR, Davis MJ, Reverter A, Ragan MA. 2012. Gene regulatory network inference: evaluation and application to ovarian cancer allows the prioritization of drug targets. Genome Med 4: 41.

Marbach D, Costello JC, Küffner R, Vega NM, Prill RJ, Camacho DM, Allison KR, DREAM5 Consortium, Kellis M, Collins JJ, et al. 2012. Wisdom of crowds for robust gene network inference. Nat Methods 9: 796–804.

Meier T, Timm M, Montani M, Wilkens L. 2021. Gene networks and transcriptional regulators associated with liver cancer development and progression. BMC Med Genomics 14: 41.

Meng J, Guan Y, Wang B, Chen L, Chen J, Zhang M, Liang C. 2022. Risk subtyping and prognostic assessment of prostate cancer based on consensus genes. Commun Biol 5: 233.

Moerman T, Aibar Santos S, Bravo González-Blas C, Simm J, Moreau Y, Aerts J, Aerts S. 2019. GRNBoost2 and Arboreto: efficient and scalable inference of gene regulatory networks. Bioinformatics 35: 2159–2161.

Morgan D, Studham M, Tjärnberg A, Weishaupt H, Swartling FJ, Nordling TEM, Sonnhammer ELL. 2020. Perturbation-based gene regulatory network inference to unravel oncogenic mechanisms. Sci Rep 10: 14149.

Mucha B, Qie S, Bajpai S, Tarallo V, Diehl JN, Tedeschi F, Zhou G, Gao Z, Flashner S, Klein-Szanto AJ, et al. 2022. Tumor suppressor mediated ubiquitylation of hnRNPK is a barrier to oncogenic translation. Nat Commun 13: 6614.

Müller S, Glaß M, Singh AK, Haase J, Bley N, Fuchs T, Lederer M, Dahl A, Huang H, Chen J, et al. 2019. IGF2BP1 promotes SRF-dependent transcription in cancer in a m6A- and miRNA-dependent manner. Nucleic Acids Res 47: 375–390.

Osma-Garcia IC, Mouysset M, Capitan-Sobrino D, Aubert Y, Turner M, Diaz-Muñoz MD. 2023. The RNA binding proteins TIA1 and TIAL1 promote Mcl1 mRNA translation to protect germinal center responses from apoptosis. Cell Mol Immunol 20: 1063–1076.

Palombo R, Verdile V, Paronetto MP. 2020. Poison-Exon Inclusion in DHX9 Reduces Its Expression and Sensitizes Ewing Sarcoma Cells to Chemotherapeutic Treatment. Cells 9. 10.3390/cells9020328.

Pauwels EKJ, Volterrani D. 2021. Coffee Consumption and Cancer Risk: An Assessment of the Health Implications Based on Recent Knowledge. Med Princ Pract 30: 401–411.

Pereira B, Billaud M, Almeida R. 2017. RNA-Binding Proteins in Cancer: Old Players and New Actors. Trends Cancer Res 3: 506–528.

Persson E, Castresana-Aguirre M, Buzzao D, Guala D, Sonnhammer ELL. 2021. FunCoup 5: Functional Association Networks in All Domains of Life, Supporting Directed Links and Tissue-Specificity. J Mol Biol 433: 166835.

Qin H, Ni H, Liu Y, Yuan Y, Xi T, Li X, Zheng L. 2020. RNA-binding proteins in tumor progression. J Hematol Oncol 13: 90.

Quattrone A, Dassi E. 2019. The Architecture of the Human RNA-Binding Protein Regulatory Network. iScience 21: 706–719.

Rawat C, Heemers HV. 2024. Alternative splicing in prostate cancer progression and therapeutic resistance. Oncogene 43: 1655–1668.

Safe S, Kothari J, Hailemariam A, Upadhyay S, Davidson LA, Chapkin RS. 2023. Health Benefits of Coffee Consumption for Cancer and Other Diseases and Mechanisms of Action. Int J Mol Sci 24. 10.3390/ijms24032706.

Sakasai R, Isono M, Wakasugi M, Hashimoto M, Sunatani Y, Matsui T, Shibata A, Matsunaga T, Iwabuchi K. 2017. Aquarius is required for proper CtIP expression and homologous recombination repair. Sci Rep 7: 13808.

Seçilmiş D, Hillerton T, Morgan D, Tjärnberg A, Nelander S, Nordling TEM, Sonnhammer ELL. 2020. Uncovering cancer gene regulation by accurate regulatory network inference from uninformative data. NPJ Syst Biol Appl 6: 37.

Seçilmiş D, Hillerton T, Tjärnberg A, Nelander S, Nordling TEM, Sonnhammer ELL. 2022. Knowledge of the perturbation design is essential for accurate gene regulatory network inference. Sci Rep 12: 16531.

Segura-Ortiz A, García-Nieto J, Aldana-Montes JF, Navas-Delgado I. 2023. GENECI: A novel evolutionary machine learning consensus-based approach for the inference of gene regulatory networks. Comput Biol Med 155: 106653.

Sequera C, Grattarola M, Holczbauer A, Dono R, Pizzimenti S, Barrera G, Wangensteen KJ, Maina F. 2022. MYC and MET cooperatively drive hepatocellular carcinoma with distinct molecular traits and vulnerabilities. Cell Death Dis 13: 994.

Shen B, Coruzzi G, Shasha D. 2023. EnsInfer: a simple ensemble approach to network inference outperforms any single method. BMC Bioinformatics 24: 114.

Shu H, Zhou J, Lian Q, Li H, Zhao D, Zeng J, Ma J. 2021. Modeling gene regulatory networks using neural network architectures. Nat Comput Sci 1: 491–501.

Silla T, Schmid M, Dou Y, Garland W, Milek M, Imami K, Johnsen D, Polak P, Andersen JS, Selbach M, et al. 2020. The human ZC3H3 and RBM26/27 proteins are critical for PAXT-mediated nuclear RNA decay. Nucleic Acids Res 48: 2518–2530.

Søndergaard JN, Sommerauer C, Atanasoai I, Hinte LC, Geng K, Guiducci G, Bräutigam L, Aouadi M, Stojic L, Barragan I, et al. 2022. CCT3-LINC00326 axis regulates hepatocarcinogenic lipid metabolism. Gut 71: 2081–2092.

Song Y-Y, Lu Y. 2015. Decision tree methods: applications for classification and prediction. Shanghai Arch Psychiatry 27: 130–135.

Subramanian A, Tamayo P, Mootha VK, Mukherjee S, Ebert BL, Gillette MA, Paulovich A, Pomeroy SL, Golub TR, Lander ES, et al. 2005. Gene set enrichment analysis: a knowledge-based approach for interpreting genome-wide expression profiles. Proc Natl Acad Sci U S A 102: 15545–15550.

Tjärnberg A, Morgan DC, Studham M, Nordling TEM, Sonnhammer ELL. 2017. GeneSPIDER - gene regulatory network inference benchmarking with controlled network and data properties. Mol Biosyst 13: 1304–1312.

Uramoto H, Izumi H, Nagatani G, Ohmori H, Nagasue N, Ise T, Yoshida T, Yasumoto K, Kohno K. 2003. Physical interaction of tumour suppressor p53/p73 with CCAAT-binding transcription factor 2 (CTF2) and differential regulation of human high-mobility group 1 (HMG1) gene expression. Biochem J 371: 301–310.

Van Nostrand EL, Pratt GA, Shishkin AA, Gelboin-Burkhart C, Fang MY, Sundararaman B, Blue SM, Nguyen TB, Surka C, Elkins K, et al. 2016. Robust transcriptome-wide discovery of RNA-binding protein binding sites with enhanced CLIP (eCLIP). Nat Methods 13: 508–514.

Wang J, Sun J, Zhang N, Yang R, Li H, Zhang Y, Chen K, Kong D. 2019. PES1 enhances proliferation and tumorigenesis in hepatocellular carcinoma via the PI3K/AKT pathway. Life Sci 219: 182–189.

Whisenant TC. 2017. Gene expression profiling of U2AF2 dependent RNA-protein interactions during CD4+ T cell activation. Genomics Data 11: 77–80.

Wu J, Maller B, Kaul R, Galabow A, Bryan A, Neuwelt A. 2024. High-Dose Acetaminophen as a Treatment for Cancer. Livers 4: 84–93.

Xie W, Qu L, Meng L, Liu C, Wu J, Shou C. 2013. PES1 regulates sensitivity of colorectal cancer cells to anticancer drugs. Biochem Biophys Res Commun 431: 460–465.

Xu C, Chen X, Zhang X, Zhao D, Dou Z, Xie X, Li H, Yang H, Li Q, Zhang H, et al. 2021. RNA-binding protein 39: a promising therapeutic target for cancer. Cell Death Discov 7: 214.

Xu J, Xu J, Liu X, Jiang J. 2022. The role of lncRNA-mediated ceRNA regulatory networks in pancreatic cancer. Cell Death Discov 8: 287.

Zeng L, Yu J, Huang T, Jia H, Dong Q, He F, Yuan W, Qin L, Li Y, Xie L. 2012. Differential combinatorial regulatory network analysis related to venous metastasis of hepatocellular carcinoma. BMC Genomics 13 Suppl 8: S14.

Zhang J, Hu K, Yang Y-Q, Wang Y, Zheng Y-F, Jin Y, Li P, Cheng L. 2020. LIN28B-AS1-IGF2BP1 binding promotes hepatocellular carcinoma cell progression. Cell Death Dis 11: 741.

Zhang Y, Qian J, Gu C, Yang Y. 2021. Alternative splicing and cancer: a systematic review. Signal Transduct Target Ther 6: 78.

Zhu G-Q, Wang Y, Wang B, Liu W-R, Dong S-S, Chen E-B, Cai J-L, Wan J-L, Du J-X, Song L-N, et al. 2022. Targeting HNRNPM Inhibits Cancer Stemness and Enhances Antitumor Immunity in Wnt-activated Hepatocellular Carcinoma. Cell Mol Gastroenterol Hepatol 13: 1413–1447.

